# A high-resolution meiotic crossover map from single-nucleus ATAC-seq reveals insights into the recombination landscape in mammalian sperm

**DOI:** 10.1101/2024.12.16.625960

**Authors:** Stevan Novakovic, Caitlin Harris, Ruijie Liu, Davis J. McCarthy, Wayne Crismani

## Abstract

Meiotic crossovers promote correct chromosome segregation and the shuffling of genetic diversity. However, the measurement of crossovers remains challenging, impeding our ability to decipher the molecular mechanisms that are necessary for their formation and regulation. Here we demonstrate a novel repurposing of the single-nucleus Assay for Transposase Accessible Chromatin with sequencing (snATAC-seq) as a simple and high-throughput method to identify and characterise meiotic crossovers from sperm nuclei. We first validate the feasibility of obtaining genome-wide coverage from snATAC-seq by using ATAC-seq on bulk haploid mouse sperm, ensuring adequate variant detection for haplotyping. Subsequently, we adapt droplet-based snATAC-seq for crossover detection, revealing over 25,000 crossovers in F1 hybrid mice. Comparison between wildtype and a hyper-recombinogenic *Fancm*-deficient mutant mouse model confirmed an increase in crossover rates in this genotype, however a distribution which was unchanged. We also find that regions with the highest rate of crossover formation are enriched for DMC1 and PRDM9, with a subset that is further enriched for DMC1 in *Fancm*-deficient mice. Our findings demonstrate the utility of snATAC-seq as a robust and scalable tool for high-throughput crossover detection, offering insights into meiotic crossover dynamics and elucidating the underlying molecular mechanisms. It is possible that the research presented here with snATAC-seq of haploid sperm could be extended into fertility-related diagnostics.

## Introduction

Meiosis is a specialised cell division which is necessary for the formation of haploid gametes in sexually reproducing species. Meiosis halves ploidy through one round of DNA replication being followed by two consecutive rounds of chromosome division. The first round of chromosome segregation (meiosis I) is a reductional division, where homologues are pulled to opposite poles. The second round of chromosome segregation (meiosis II) is an equational division where sister chromatids are pulled to opposite poles (Hunter, 2015). At the onset of meiosis, hundreds of DNA double-strand breaks (DSBs) are formed (Lam & Keeney, 2015). Homologous recombination is used to repair the DSBs as either crossovers (COs) or non-crossovers (NCO) (Gray & Cohen, 2016; Hunter, 2015; Szostak et al., 1983). Meiotic crossovers are large reciprocal exchanges between homologous chromosomes. For chromosomes to segregate correctly at the first meiotic division, all homologue pairs must have at least one “obligate” crossover. Loss of the obligate crossover leads to aneuploid gametes and a reduction in fertility (Hunter, 2015). Aneuploid gametes used in fertilisation can lead to aneuploid karyotypes, such as trisomy 21 in humans (Hassold & Sherman, 2000; Sherman et al., 1991; Warren et al., 1987). The obligate crossover is linked to a phenomenon known as crossover interference, where the occurrence of one chromosome reduces the probability of another crossover on the same chromosome in the same meiosis in a distance dependent manner (Berchowitz & Copenhaver, 2010; Hunter, 2015).

Assaying for genome-wide crossovers remains challenging, despite more than a century since the discovery of meiotic crossing over in drosophila and how it informs on physical chromosome structure and enables genetic map construction (Morgan, 1910; Sturtevant, 1913). These detection challenges mostly relate to the need for generating and genotyping pedigrees that require at least three generations and many offspring.

Reporter assays continue to play an essential role in deciphering the mechanisms of double- strand break repair and crossover formation. Some examples include DSB hotspot analysis, linked phenotypic markers and tetrad analysis across an array of species (Cole et al., 2014; Fogel & Mortimer, 1971; Francis et al., 2007; McClintock, 1931; Morgan, 1910; Schwacha & Kleckner, 1994; Yelina et al., 2013). These assays can allow quantitative analysis of a variety of parameters at a single locus or adjacent loci e.g. formation of DSBs, crossovers, non- crossovers and, repair on sister chromatids. A strength of their methods is the reproducibility once established and potential for comparisons across experimental conditions. On the other hand, cytological analysis of DSB and crossover formation inform on chromosome-scale and genome-wide events. However, these methods, while providing rich and important data tend to be more labour intense, particularly for quantification of events being scored.

Whole-genome single-gamete sequencing provides the opportunity to combine the advantages of different styles of assays for crossover detection and quantification. Single-gamete sequencing for genome-wide crossover analysis can reduce the number of generations required to access the recombined products of meiosis. Interest in understanding crossover regulation has driven the adaptation of various technologies to sequence individual gametes (Lu et al., 2012; Bell et al., 2019; Hinch et al., 2019; Luo et al., 2019; Sun et al., 2019; J. Campoy et al., 2020; Tsui, et al., 2023; Xie et al., 2023) and the development of methods for analysis of the data (Carioscia et al., 2022; Lyu et al., 2022). Notably, droplet-based gamete sequencing methods are a powerful tool for investigating genome-wide crossover formation, because despite the low level of coverage typically obtained per cell, the coverage is sufficient to determine haplotypes (J. Campoy et al., 2020; Leung et al., 2021; Lyu et al., 2022; Tsui, et al., 2023). However, despite advances in single gamete sequencing approaches, specialised kits used in previous studies have been discontinued which results in established methods becoming unavailable. We therefore sought to establish a method that we consider should remain accessible for longer or could be re-established with in-house reagents in the case of product discontinuation. The Assay for Transposase Accessible Chromatin using sequencing (ATAC-seq) was developed for profiling chromatin accessibility at a genome-wide scale (Buenrostro et al., 2013). This method of DNA library preparation could be adapted for single-nucleus sperm sequencing for crossover detection applications, which we predict as it can generate reads genome-wide – in some cell lines and tissues – and has already been used for single cell sequencing assays (Cusanovich et al., 2015). Further, open-source methods for Tn5 expression and purification are well established which can facilitate in-house development of ATAC-based methods. However, it is less clear the extent to which gametes can be sequenced with ATAC-based methods given the different chromatin structure – which is protamine rich – in mammalian sperm compared to somatic tissue.

We demonstrate here that snATAC-seq libraries can be used to sequence genomic DNA for crossover quantification and analyses. Using hybrid mice, from a cross between the two strains C57BL/6J and FVB, we isolated haploid sperm and prepared libraries using modified snATAC-seq methods. We obtained high quality libraries which had ample coverage for variant calling, phasing and crossover detection. Further the number of high-quality cells recovered was far greater than other methods that we have used. Using an established hyper- crossover mouse model, we could also demonstrate that this robust method can detect changes in crossover frequencies, opening scope for other applications in research and diagnostics.

## Results

### Bulk ATAC-sequencing libraries using sperm nuclei produce genome-wide coverage

ATAC-seq assays use the enzymatic activity of Tn5 to generate DNA libraries for high throughput sequencing. Tn5 simultaneously cuts accessible regions of double stranded DNA while ligating sequencing adapters compatible with high-throughput whole-genome sequencing platforms (Adey et al., 2010; Buenrostro et al., 2013; Goryshin & Reznikoff, 1998). However, sperm chromatin is particularly condensed and protamine rich (Hammoud et al., 2009; Oliva, 2006), and it was unclear if libraries could generate read coverage suitable for haplotyping and CO measurement. Therefore, we first sought to test if sufficient coverage of sperm DNA could be obtained in bulk ATAC-seq samples to justify using ATAC-seq with single-sperm nuclei.

We first performed ATAC-sequencing by isolating 50,000 bulk-sorted haploid sperm nuclei extracted from F1 (C57BL/6J x FVB/N) hybrid mice, using our SSNIP-seq protocol (Novakovic et al., 2022) for high quality sperm nuclei isolation. We expressed and purified Tn5 (Supplementary Figure 1) using a plasmid previously described (Picelli et al., 2014) and then generated tagmented sequencing libraries using our home-made and commercially available hyperactive Tn5 enzymes. Our sequencing libraries served as a proof-of-concept, demonstrating genome wide sequencing coverage derived from sperm nuclei is attainable via ATAC-sequencing.

Our bulk ATAC-sequencing libraries were used to generate approximately 152 to 189 million paired-end reads, with an average coverage of approximately 6 to 7x across the mouse genome (Table 1). Our libraries exhibited minimal GC-bias and covered 42-50% of the informative SNPs between C57BL/6J and FVB backgrounds (Table 1), which we used for haplotyping and crossover detection previously (Tsui et al., 2023). While our ATAC-sequencing data does not map homogenously throughout the genome at nucleotide resolution (Table 1, Figure 1A-C) –as is expected with ATAC-seq – with the high number of SNPs in the genome we predicted it would be sufficient for haplotyping and crossover calling. Our prediction was based on the observation that when considering “bins” throughout the genome (Figure 1A-C), reads map relatively homogenously to broader regions, even if at the resolution of a gene there is skewing of reads towards regulatory elements and exons (Figure 1C). We therefore proceeded to test and repurpose single-gamete ATAC-sequencing methods for meiotic crossover detection.

**Figure 1:**
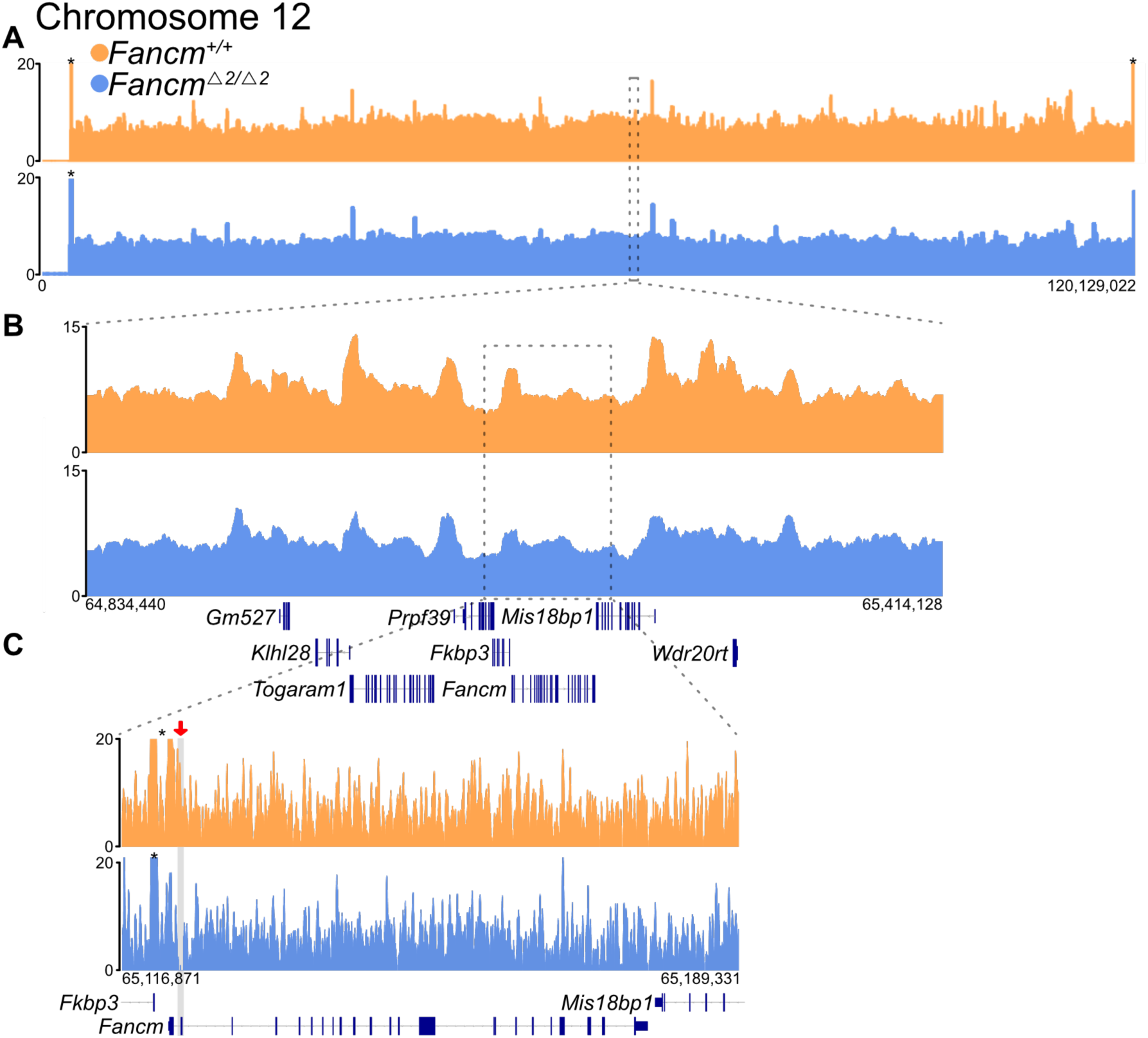
Bulk sequencing of tagmented sperm nuclei generates fragments that map broadly with genome-wide coverage. Library were generated by Tn5 fragmentation of haploid sperm nuclei from wild type and mutant F1 hybrid mouse testis, with reads aligned to the mouse reference genome (mm39). Tracks show reads from wildtype (orange) and mutant (blue) samples, generated with home- made Tn5, aligned to: A) The length of chromosome 12 (64 kb bins). B) A zoomed-in ∼580 kb region (chr12: 64,834,440 to 65,414,128; 10 kb bins). C) A zoomed-in ∼72 kb region (chr12: 65,116,871 to 65,189,331; 100 bp bins). The grey box highlights the locus of exon 2 of *Fancm;* indicated by a red arrow for clarity. *Maximum displayed coverage threshold set to 20.

**Table 1.** Bulk-sequencing results from sperm nuclear DNA prepared with Tn5. Sequence output metrics are provided for bulk ATAC-seq libraries produced from haploid sperm DNA.

| Tn5 source | Genotype | Total Mapped read pairs | Read length (bp) | Average genome-wide coverage | %GC | Proportion of informative SNPs with a least one mapped read* |
| --- | --- | --- | --- | --- | --- | --- |
| Home-made | *Fancm*<sup>+/+</sup> | 188,998,455 | 150 | 6.7 | 49 | 49.7% |
| Commercial | *Fancm*<sup>+/+</sup> | 151,991,673 | 150 | 5.5 | 50 | 41.5% |
| Home-made | *Fancm*<sup>Δ2/Δ2</sup> | 179,633,072 | 150 | 6.1 | 51 | 45.4% |
| Commercial | *Fancm*<sup>Δ2/Δ2</sup> | 162,885,127 | 150 | 5.8 | 51 | 41.5% |

### snATAC-seq library generation, pre-processing and filtering

To assay meiotic crossovers with high throughput, we generated single-sperm ATAC-sequencing libraries (10x Genomics) using FACS-sorted haploid spermatocytes isolated from six F1 hybrid male mice; three F1.*Fancm^+/+^* and three F1.*Fancm*^112/112^. Coverage estimates of the snATAC data were performed through assessing the number of fragments at common regions of open chromatin across samples. Read coverage showed that the snATACseq data from haploid sperm had relatively homogeneous coverage across bins of the genome (Figure 2A), replicating the genome-wide coverage observed in the bulk ATAC-seq data (Figure 1; Table 1).

**Figure 2:**
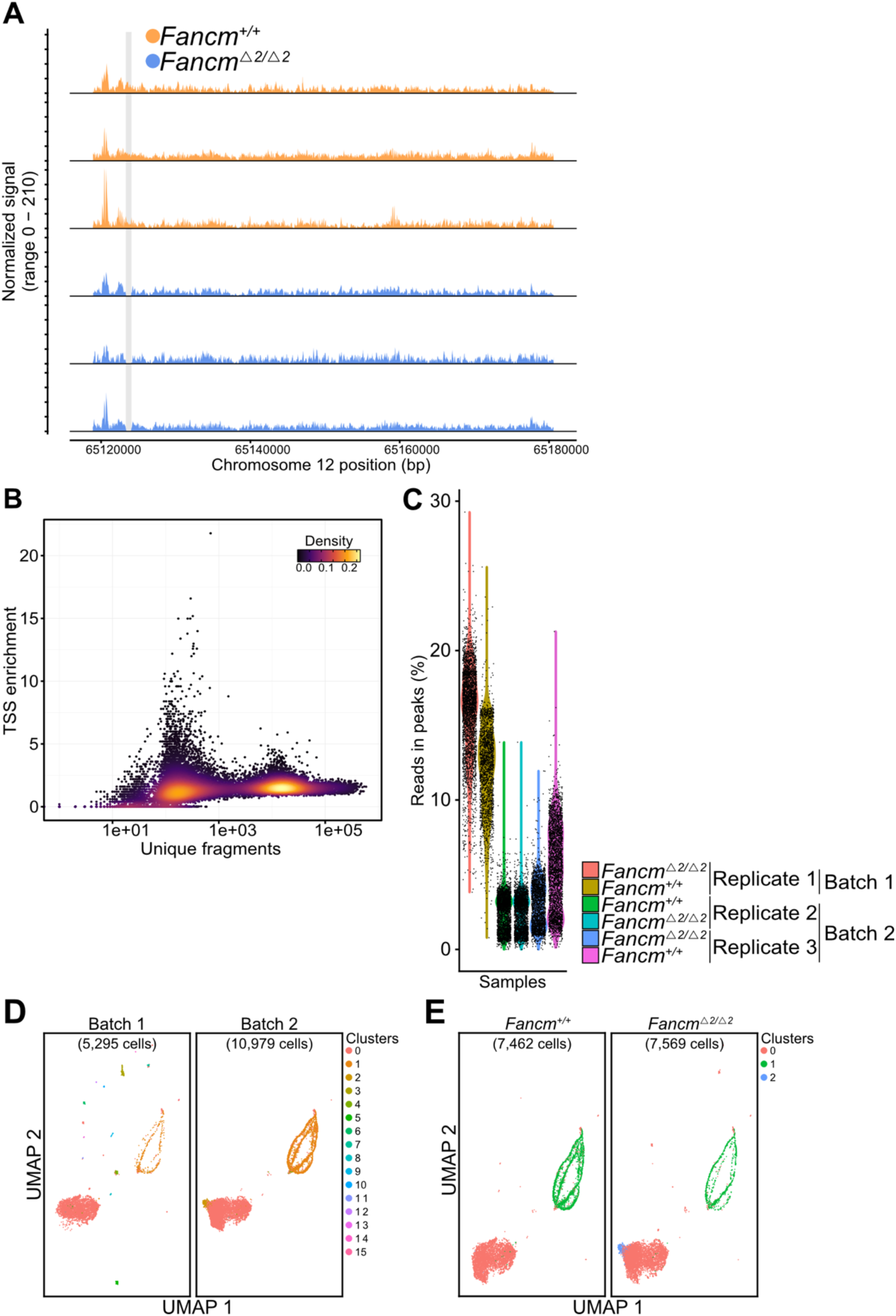
Quality control and validation of the haploid sperm snATAC-seq libraries. A) Comparison of genome coverage across a ∼60 kb window of chromosome 12 (Chr12: 65,120,000 to 65,180,000) from each mouse single gamete library. The locus contains *Fancm*, with no reads mapping to the exon 2 of the F1.*Fancm*^/12//12^, which carry biallelic deletions (grey) of exon 2. B) scatter plot of snATAC-seq data displaying quality control metrics (unique fragments per cell and TSS enrichmint). Low TSS enrichment highlights the condensed nature of sperm chromatin. C) Violin plot showing the percentage of reads that cluster into peaks for each wild type and mutant replicate. Three replicates (wild-type / mutant pairs) were sequenced across two sequencing runs (batch 1 and 2). D) UMAP visualisations of snATAC-seq of haploid sperm, from sequencing batches 1 and 2, showing cells grouped in 9 clusters. E) UMAP of snATAC data for wildtype and mutant samples, post-filtering.

Given that sperm chromatin is more condensed than in other cell types, bioinformatic tools Signac (v1.13.0) (Stuart et al., 2021) and Seurat (v5.0.3) (Stuart et al., 2019) were employed to assess chromatin accessibility and evaluate the quality of the dataset. We calculated the number of fragments per cell, which provided a measure of chromatin accessibility for each cell in the sample group. Additionally, Transcription Start Site (TSS) enrichment scores were computed to indicate the accessibility of chromatin near TSS regions, reflecting the regulatory activity in these areas. These metrics were visualized using a density scatter plot (Figure 2B). The relative number of counts per cell aligns with the known inaccessibility of sperm chromatin (Oliva, 2006). snATAC-seq libraries were produced from three biological replicates of each genotype (F1.*Fancm^+/+^* and F1.*Fancm*^112/112^) in two batches and sequenced in multiple runs (Figure 2C-E). When assessing the percentage of reads in peaks (Figure 2C), the data segregated into two groups, indicating possible batch effects, however this does not affect the ability to detect crossovers (below). When exploring the overall structure of the data in a UMAP plot, the samples from the first sequencing batch appeared to have an increased number of artefacts, indicated by the large number of smaller clusters consisting of a few cells (Figure 2D). The artefacts were removed, and the top genetic markers were identified from each cluster, for each genotype. When comparing wild-type and mutant samples from the filtered data set, gene set enrichment analysis (see Methods) indicated that cluster 0 corresponds to late-stage spermatocytes, while cluster 1 represents early-stage spermatocytes (Figure 2E). Cluster 2 could not be identified, indicating it may consist of contamination, unfiltered artefacts, or an unannotated cell state. However, without single-cell RNA-sequencing or cytological analysis, confirming the exact identify of this cluster remains challenging. Despite this ambiguity, the results confirm the correct cell types were sequenced with minimal contamination, making the data suitable for further processing and crossover calling analysis.

The single sperm sequencing library was filtered for barcodes which were present with >10k high quality fragments and excluded likely doublets, and cells with an incomplete complement of chromosomes. The final dataset included more than 3800 haploid sperm genomes from *Fancm^+/+^* and *Fancm*^112/112^ samples (Table 2, Supplementary table 1). Each cell was sequenced to an average depth of 0.01x of the haploid genome, covering approximately 30 million base pairs per cell on average. About 0.65% of the heterozygous SNPs between the two mouse strains had at least one mapped read, which is sufficient for haplotyping (Table 2). To assess alignment bias and sequencing bias we analysed allelic segregation ratios and found that throughout the genome most loci segregated very close to a ratio of 1:1, in the three F1.*Fancm^+/+^* and three F1.*Fancm*^112/112^ samples (Figure 3). Some telomeric skewing remains on a limited number of chromosomes after filtering repetitive regions, typically representing a mapping bias towards the reference genome.

**Figure 3.**
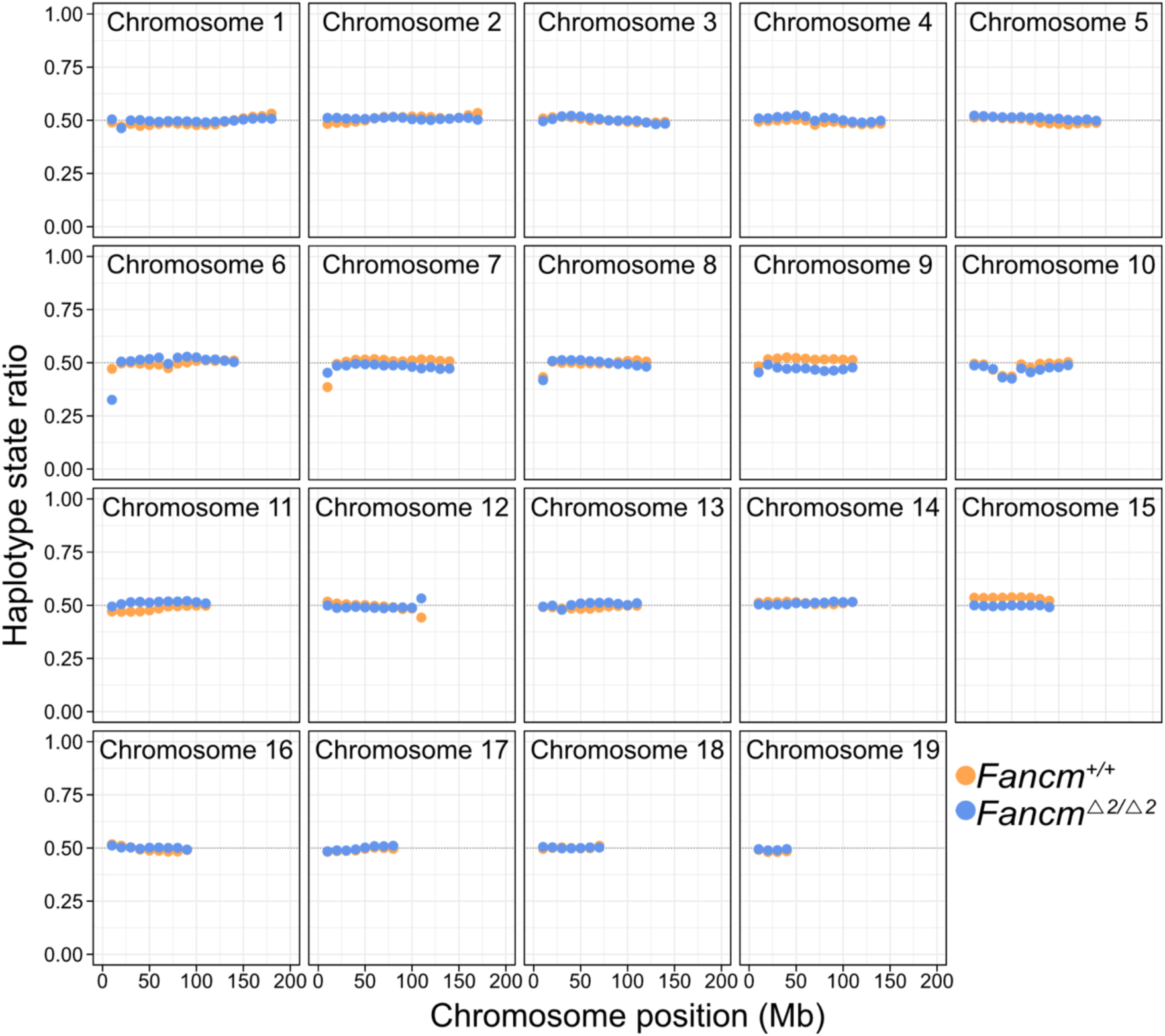
Marker segregation for the 19 autosomal chromosomes in *Fancm* wild type and mutant snATAC-seq data. Genome-wide patterns of marker segregation from gametes produced by F1(C57BL/6J x FVB) mouse with two haplotypes were calculated in chromosome bins (of size 10 Mb) and found to match Mendelian segregation expectations, except for sub- telomeric regions (excluded from analysis due to mapping biases in repetitive regions). Hypothesis testing using a binomial test was performed to evaluate if marker segregation ratios differ from 0.5; no significant differences were observed in any chromosomes. The y-axis units of haplotype state ratio represent FVB or C57BL/6J ratios for given genomic bins, “1” represents the alternate allele, which is FVB, and “0” represents the reference allele C57BL/6J. Most genomic regions have a haplotype state ratio close to 0.5, which is consistent with chromosome segregation showing expected Mendelian ratios.

**Table 2:** Summary of crossovers detected from ATAC-sequencing of single sperm. Data summarized from ATAC-sequencing results. Total numbers of COs from *Fancm*^+/+^ and *Fancm^Δ2/Δ2^* samples were broken down into chromosomes with single, double, and >2 COs. *Individual gametes were identified and filtered for a unique barcode which was represented by >10 k high quality fragments (see Methods).

| Genotype | Cells* | Single crossovers | Double crossovers | >2 crossovers | Total crossovers |
| --- | --- | --- | --- | --- | --- |
| *Fancm*<sup>+/+</sup> | 2,251 | 14,778 | 1,402 | 63 | 16,243 |
| *Fancm*<sup>Δ2/Δ2</sup> | 1,581 | 10,496 | 1,617 | 104 | 12,217 |

### Meiotic crossover detection with snATAC-seq libraries

Using sgcocaller and comapr (Lyu et al., 2022), we identified >25,000 meiotic crossovers within our combined snATAC-sequencing samples. We found that crossover detection was robust and was not affected by coverage within the ranges of read depth in our samples (Figure 4).

**Figure 4:**
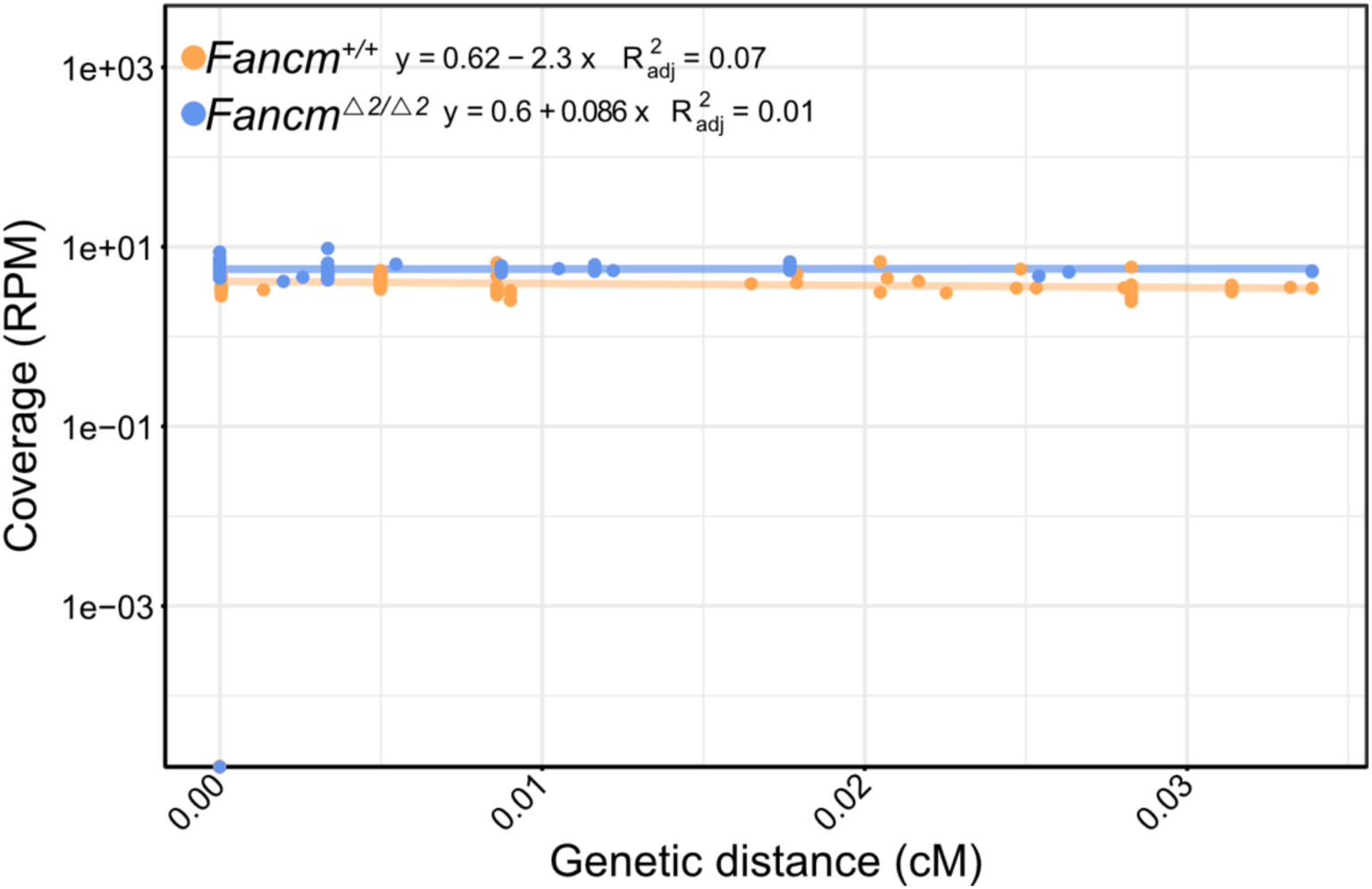
Analysis of genetic distance (cM) as a function of coverage reveals a robust assay for crossover detection. Coverage is defined as reads per million [RPM] mapped reads per sample. We find no significant correlation between genetic distance and coverage for both wildtype and mutant F1 mouse samples using snATAC-seq, in 10kb bins.

Notably, the average number of crossovers inferred per sperm were higher in F1.*Fancm*^/12//12^ samples compared to F1.*Fancm^+/+^* controls, with a median of 12 and 10 crossovers identified per sperm, respectively (Two-sided Wilcoxon signed rank test with continuity correction, *p < 2.2*×*10^-16^*; Figure 5A), and the genetic map lengths were significantly longer in mutant (1,172.4 cM) than wild type (1,015.2 cM) samples (Two-sided Wilcoxon signed rank test with continuity correction, *p < 2.2*×*10^-16^*; Figure 5B). These findings align with our previous work indicating *Fancm* acts as a meiotic crossover suppressor in mammals (Tsui et al., 2023). Most individual chromosomes had genetic lengths of at least 50 cM, indicating good variant detection along the length of each chromosome (Figure 5C). The differences in genetic distance found in *Fancm* mice were observed along the length of individual chromosomes in bins of 10 Mb (Supplementary table 2). Crossover numbers were elevated in mutant mice at a chromosome level, with adjusted p-values above 0.01 in all chromosomes except 4, 5, 9, 15 and 19, as shown by both Kruskal-Wallis and permutation testing with false discovery rate (FDR) correction (Supplementary table 2). Although each chromosome had increased crossovers in the mutant mice, the general crossover distribution – at any given locus – remained comparable between both genotypes, with no significant difference between the 10 kb bins (Permutation testing with FDR correction, *p > 0.05*; Figure 5D). These results are consistent with our previous findings using single-sperm, bulk sequencing, and PCR-based methods for crossover detection (Tsui et al., 2023), suggesting that ATAC-seq provides a robust tool for high-throughput crossover detection.

**Figure 5:**
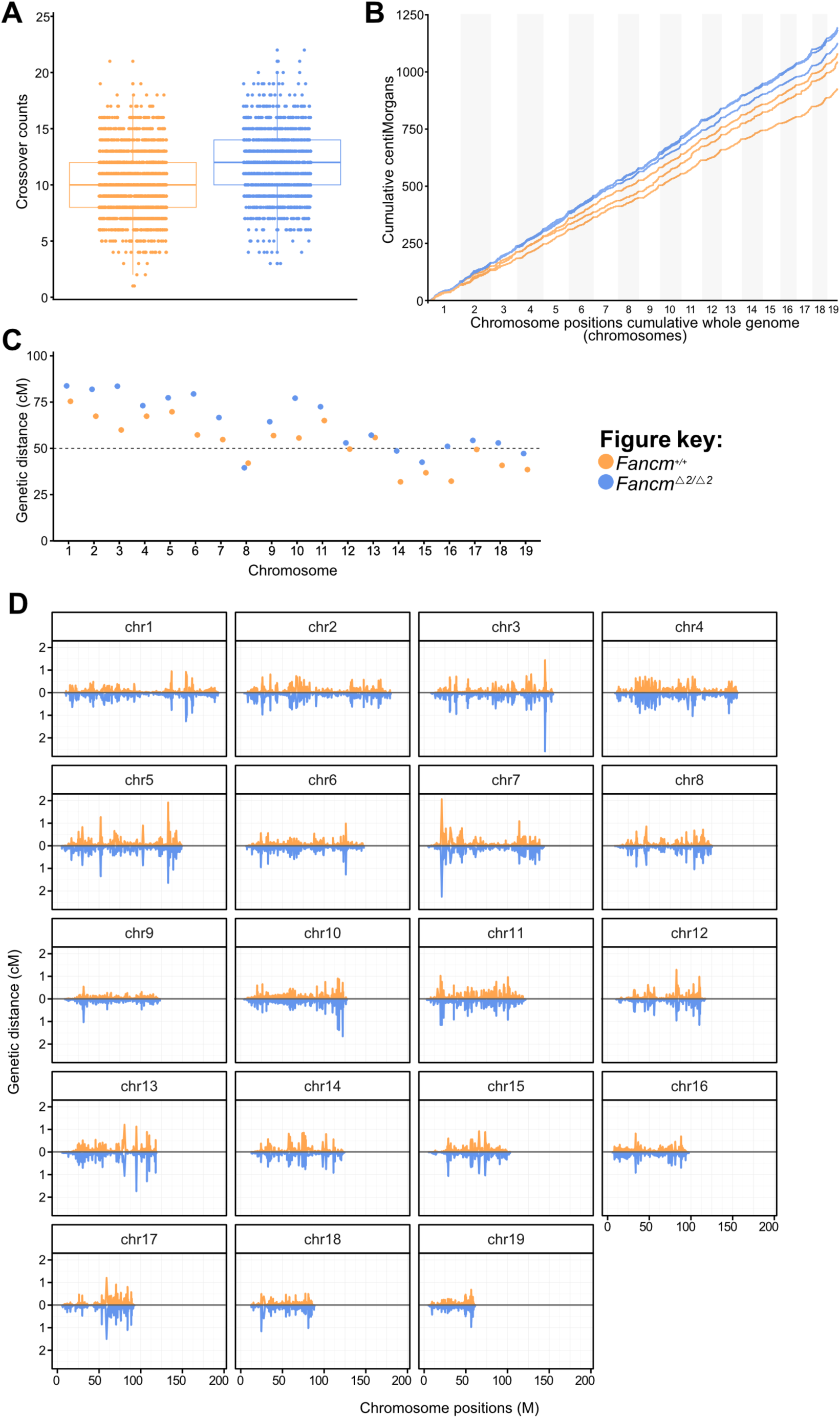
snATAC-seq profiling of meiotic crossovers. Crossovers were assayed using snATAC-sequencing of isolated haploid mouse sperm. A) Distribution of CO frequency assayed per haploid sperm (n = 3 animals per genotype). Boxplot represents median, upper, and lower quartile; whiskers represent the lowest and highest values within the 1.5 interquartile range. B) Recombination rates measured for F1.*Fancm^+/+^* and F1.*Fancm^/12//12^* samples. Observed crossover fractions were converted into genetic distances (centiMorgans) via the Kosambi mapping function and presented as cumulative centiMorgans across the genome. C) Average recombination frequency (cM) per chromosome per mutant and wild type samples. D) Recombination rates (in cM) measured per 10 kb window along each chromosome position (M, megabases) for *Fancm^+/+^* (top) and *Fancm^/12//12^* (bottom, flipped) autosomes reveal the increased crossover rate in mutant mice.

### Genome-wide crossover distributions detected with snATAC-seq

Additionally, we investigated whether crossover interference could be detected with the snATAC-seq method of crossover detection. Permutation testing via label swapping was used to generate random null distributions of inter-crossover distances to simulate the absence of crossover interference (null hypothesis), as we conducted previously (Tsui et al., 2023) . We next filtered our dataset to chromosomes with only two crossovers (1,402 wild type and 1,617 mutant chromosomes; Table 2) and calculated the inter-crossover distance for each pair of crossovers. Analysis of the spacing between two crossover events on the same chromosome revealed that inter-crossover distance did not fit a random distribution in both wild type and mutant samples, indicative of a functioning crossover interference mechanism (Figure 6A-C). The observed median inter-crossover distance was 88.8 Mb in F1.*Fancm*^+/+^, and reduced to 82.6 Mb in F1.*Fancm^/12//12^*; pairwise comparisons using Wilcoxon rank sum test with continuity correction, *p < 2***×***10^-7^* (Figure 6A-C). The increase in crossovers in the absence of *Fancm* is consistent with previous findings (Blary et al., 2018; Crismani et al., 2012; Fernandes et al., 2017; Girard et al., 2014, 2015; Lorenz et al., 2012; Mieulet et al., 2018; Séguéla-Arnaud et al., 2015, 2016; Tsui et al., 2023), suggesting the additional crossovers generated in gametes lacking *Fancm* are likely to arise from the class II (non-interfering) crossover pathway. These findings suggest that snATAC-seq libraries generated with methods here and using SSNIP-seq (Novakovic et al., 2022) can be used for high-throughput and genome-wide measurements of crossover interference.

**Figure 6:**
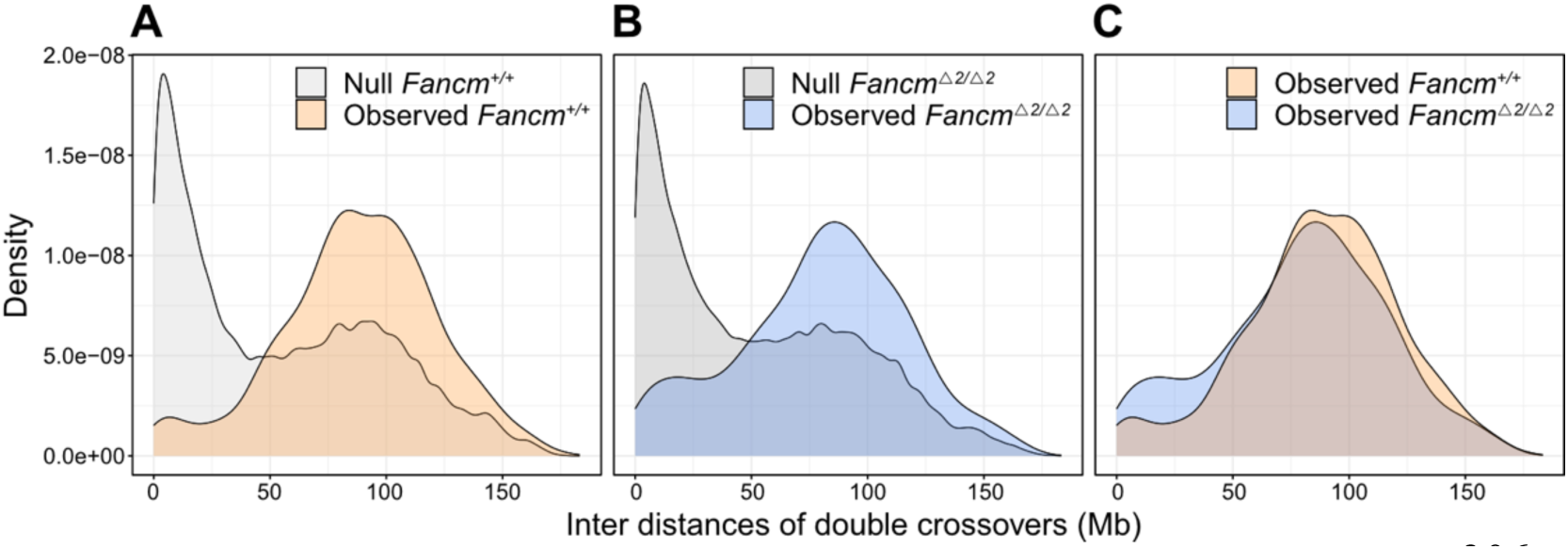
Analysis of crossover interference in haploid sperm using single nuclei ATAC-seq data. The distance of observed crossovers was compared to the null hypothesis generated from permutation via label swapping to simulate the absence of crossover interference. A) Median distances for observed double crossover chromosomes from wild type samples was approximately 88.8 Mb, compared to 0.024 Mb in the null hypothesis. Pairwise comparisons using Wilcoxon rank sum test with continuity correction*, p = 2×10^-16^*. B) Median distances for observed double crossover chromosomes from mutant samples was approximately 82.6 Mb, compared to 0.0103 Mb in the null hypothesis. Pairwise comparisons using Wilcoxon rank sum test with continuity correction, *p < 2×10^-16^*. C). Median inter-crossover distances were reduced by 6.2 Mb in mutant crossovers, with an interquartile range of 5.9 Mb. These findings suggest the additional crossovers in *Fancm*-deficient mice are likely to be derived from the non-interfering (type II) CO pathway. Pairwise comparisons using Wilcoxon rank sum test with continuity correction, *p < 2×10^-7^*. Non-parametric statistical testing was used due to non-normal data distribution.

### Crossover hotspots are associated with PRDM9 and DMC1 recombination hotspots

Crossovers identified in the snATAC-seq dataset here correlate with published PRDM9 binding locations, identified through ChIP-seq analysis from C57BL/6 mice (Biot et al., 2024). Permutation testing was conducted by comparing the mean signal at overlapping regions with the null distribution, generated by shuffling COs. P-values were calculated by comparing the observed mean signal to the null distribution, testing if the association differs from a random distribution. The results indicated there is a significant association between the COs, pooled from both genotypes, and PRDM9 binding locations (*p* < *1×10^-4^*; Figure 7A), indicating a broad requirement for PRDM9 for crossover localisation. COs from both genotypes were pooled as no differences in distributions were detected (Fig. 5D), even if there are generally more COs in *Fancm-*deficient mice. However, there was no association between the COs (this study) and C57BL/6 DMC1 binding sites (Biot et al., 2024), (permutation test, *p = 0.3*; Figure 7B).

**Figure 7:**
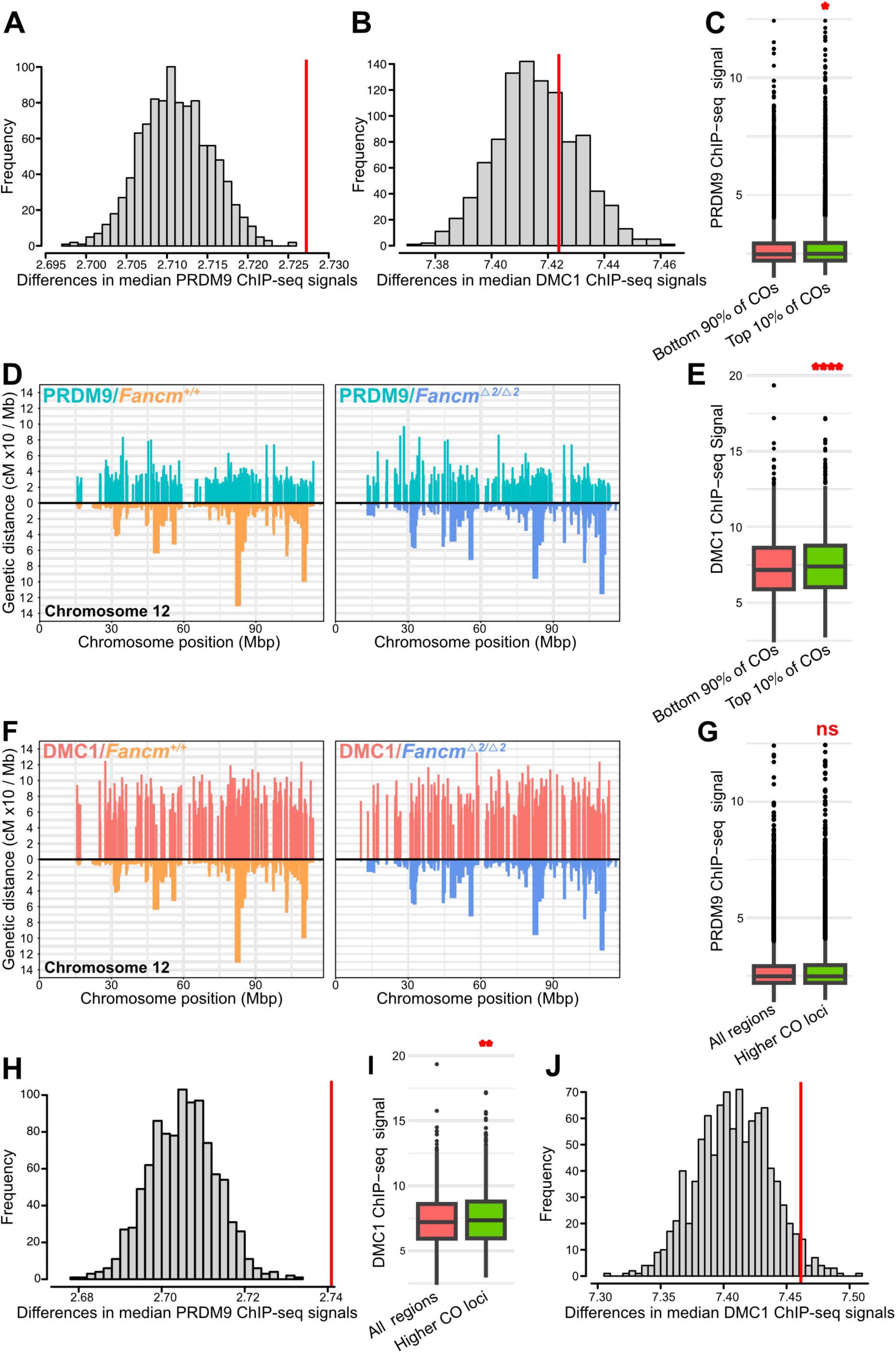
Crossover sites in (FVBxC57BL/6J) have a strong association with PRDM9 binding sites, and to a lesser extent. A) Permutation test to compare observed means with null distribution of PRDM9 ChIP-seq with all COs identified in snATAC-seq samples, *p < 1×10^-4^*. B) Permutation test to compare observed means with null distribution of DMC1 ChIP- seq with all COs identified in snATAC-seq samples; *p = 0.3*. C) One-sided Wilcoxon signed rank test for PRDM9 ChIP-seq signals compared to the top 10% of COs identified in snATAC- seq; *p = 0.02*. D) Visual representation of the top 10% of COs and overlapping PRDM9 ChIP- seq peaks. E) One-sided Wilcoxon signed rank test DMC1 ChIP-seq signals compared to the top 10% of COs (genetic distance multiplied by 10 for plotting) identified in snATAC-seq; *p < 1.5×10^-7^*. F) Visual representation of the top 10% of COs (genetic distance multiplied by 10 for plotting) and overlapping DMC1 ChIP-seq peaks. G) One-sided Wilcoxon signed rank test for PRDM9 ChIP-seq signals compared to the additional COs identified in F1.*Fancm^/12//12^* samples, compared to all other COs; *p = 0.14*. H) Permutation test to compare observed means with null distribution of PRDM9 ChIP-seq with additional F1.*Fancm^/12//12^* COs; *p < 1×10^-4^*. I) One-sided Wilcoxon signed rank test for DMC1 ChIP-seq signals compared to the additional COs identified in F1.*Fancm^/12//12^* samples, compared to all other COs; *p = 0.002*. J) Permutation test to compare observed means with null distribution of DMC1 ChIP-seq with additional F1.*Fancm^/12//12^* COs; *p = 0.03*.

COs hotspots were identified by sub-setting the data for the top 10% of COs in both F1.*Fancm^/12//12^* and F1.*Fancm*^+/+^ genotypes. Hotspots were enriched for PRDM9 ChIP-seq signals; one-sided Wilcoxon signed rank test with continuity correction, *p = 0.02* (Figure 7C- D). There was a strong, significant, association between the hotspots and DMC1 ChIP-seq signal (one-sided Wilcoxon signed rank test with continuity correction, *p = 1.5×10^-7^*; Figure 7E-F). However, there were no significant differences in either DMC1 or PRDM9 peak association with CO sites when considering genotype, F1.*Fancm^/12//12^* and F1.*Fancm*^+/+^, as a variable.

The F1.*Fancm^/12//12^* mutants have an increased number of COs, which are consistent with type II CO/NCO distributions (Figure 6A-C). When assessing the ChIP-seq signal in the loci with increased crossover frequencies in F1.*Fancm^/12//12^* compared to all F1.*Fancm*^+/+^, there was not a significant association with PRDM9 crossovers (Figure 7G-H). Permutation testing, however, indicated that there was a significant association between the extra crossovers, compared to the null distribution. This is likely due to the strong association with PRDM9 and all COs, observed in Figure 7A. However, DMC1 showed enrichment for the ‘extra’ COs identified in F1.*Fancm^/12//12^* compared to F1.*Fancm*^+/+^ (one-sided Wilcoxon signed rank test with continuity correction, *p = 0.02*, which was confirmed by permutation testing, *p = 0.03*’; Figure 7I-J).

## Discussion

### Tn5-based single gamete library preparation can be used for sequencing library production

The detection of meiotic crossover events has historically relied on pedigree data, large population cohorts, or cytogenetics. However, single-gamete library sequencing methods allow one individual to serve as the exclusive source of data for high-throughput and accurate crossover detection. These single-gamete approaches are scalable with several studies having developed methods for library preparation and crossover calling software (Bell et al., 2019; Campoy et al., 2020; Carioscia et al., 2022; Hinch et al., 2019; Lu et al., 2012; Lyu et al., 2022; Sun et al., 2019; Tsui et al., 2023). Despite these advantages, there are some practical risks than can limit the uptake of single-gamete sequencing for investigating crossover regulation, which we have aimed to partially mitigate in this study: some published methods employ reagents that are no longer commercially available (Bell et al., 2019; Tsui et al., 2023). Therefore, we sought to demonstrate proof-of-concept for single gamete library preparation with a view to then use this data for crossover detection. We also suggest that it is important to be able to adapt methods with home-made reagents to protect against flux of commercial kits. There is precedent for such an approach, e.g., plate-based methods have been established for library generation with somatic cells where bulk-tagmented nuclei are sorted into individual wells of 384-well plate for unique barcoding (Xu et al., 2021). We anticipate that single-gamete sequencing with ATAC-seq is amenable to plate-based approaches, using home-made Tn5 (Picelli et al., 2014), allowing for a cheaper alternative to commercial approaches for a smaller number of samples, typically in the hundreds.

### snATAC adaption for crossover calling from sperm

Here, we repurposed a single-nuclei ATAC-library preparation and sequencing method as a straightforward, reproducible, and highly scalable approach for genome-scale haplotyping of haploid sperm genomes. For sample preparation, we used our previously published SSNIP-seq nuclei isolation protocol (Novakovic et al., 2022), which allows for isolation of high-quality nuclei suitable as input for ATAC-sequencing. The library preparation assay itself can be completed in a single day with the collection of the sample, preparation to library generation and amplification and QC. We sequenced the genomes of 3,842 haploid mouse sperm, achieving uniform coverage across the genome at the kilobase scale. Using our established bioinformatic approaches (Lyu et al., 2022) we analysed the sequencing data to detect over 30,000 crossovers. We validated the utility of this novel approach by using our established mouse model, F1.*Fancm^/12//12^*, which has increased crossover rates (Tsui et al., 2023), whereby we showed that the snATAC-seq method detects this altered crossover behaviour. We also showed that they assay is highly robust for crossover detection, as there was no observed correlation between coverage and crossover rates, within the range of coverage that we obtained. Further, with the high number of nuclei that were sequenced, this study offers the best resolution to date of the effect of loss of function of *Fancm* in mammals, compared to our previous work. We show that there is a genome-wide increase in crossover rates in the absence of *Fancm*, these crossovers occur in the same loci as in the wildtype, and the effect in the mutant appears to be an increase in amplitude of what occurs in the wildtype. This suggests that the same DSB formation mechanisms are used, but the fate of some DSB repair intermediates progress down a crossover formation rather than non-crossover pathway.

The association between CO hotspots, PRDM9 and DMC1 binding has previously been established (Baudat et al., 2010; Hinch et al., 2019; R. Li et al., 2019; Myers et al., 2010; Parvanov et al., 2010). It is therefore not unexpected that the COs observed in this study show a similar association, however it is important to note the degree of shared crossover hotspots in our data (Figure 7D, 7F) with the binding sites of pro-crossover factors from independent work (Biot et al., 2024). The loci that account for the additional type II COs observed in *Fancm-* deficient mice are located in regions enriched for DMC1, indicating they are repaired by canonical crossover pathways. Although there was no specific enrichment for PRDM9 at these COs, compared to other COs, the strong association with PRDM9 and the complete dataset of COs might reduce the sensitivity to detect a further increase of PRDM9 at these extra crossovers in *Fancm*-deficient mice. DMC1 is only enriched in regions containing high levels of COs, and it is currently unclear why this is. It may be that it is simply easier to detect an increase in DMC1signal from its baseline levels. Another possibility is that increases in DMC1 levels, when pushed upwards, can further increase crossover rates. However, further experiments in future studies would be required to test these hypotheses.

## Conclusion and future perspectives

Our development of snATAC-seq for F1 sperm, coupled with bioinformatic analysis, establishes this as a reliable method for studying crossover events and the mechanisms required for their formation. This approach has revealed that *Fancm* plays a key role in limiting the formation of class II crossovers mediated by intrinsic levels of DMC1. We anticipate these findings and tools will have broader applications in fertility research and diagnostics, potentially aiding in the quantification of sperm aneuploidy linked to chromosome segregation errors (Templado et al., 2013). Future studies could further examine the interaction of *Fancm* with other crossover pathways, advancing our understanding of recombination and reproductive health.

## Acknowledgements

We are grateful to all members of the Crismani laboratory for helpful comments on the manuscript. Thank you to Tim Semple for helpful discussions about ATAC-seq methods and to Adam Thomas for assistance with Tn5 production. WC and DJM receive Fellowships and funding related to this work from the Australian National Health and Medical Research Council (GNT1129757, GNT1112681, GNT1185387).

## Authors’ contributions

All authors wrote, reviewed and discussed the manuscript. SN and CH prepared the figures. All authors read and approved the final manuscript.

## Ethics statement

All experimental procedures were approved by the St. Vincent’s Hospital Melbourne Animal Ethics Committee.

## Methods

### Animals

All animal experiments were approved by the Animal Ethics Committees at St Vincent’s Hospital Melbourne and conducted in accordance with Australian NHMRC Guidelines on Ethics in Animal Experimentation. All mice were housed at the BioResources Centre, St. Vincent’s Hospital, in a controlled environment with a 12-hour light/dark cycle, and with food and water provided *ad libitum*. The *Fancm^Δ2/Δ2^* mice used in this study were previously described (Tsui, et al., 2023).

### Isolation of haploid nuclei

Haploid nuclei were isolated from testicular single-nucleus suspensions and enriched for haploids using FACS as previously described (Novakovic et al., 2022). Briefly, testes were dissected out from adult mice and placed in 1 mL of chilled Nuclei EZ Lysis Buffer (Sigma). The testes were gently squeezed with pointed tweezers to releases seminiferous tubules into the solution. Simultaneously, the spleen was also dissected and homogenised with 2 mL Dulbecco’s Phosphate Buffered Saline (DBPS) before being passed through a 40 μm strainer; this sample served as a diploid control for flow cytometry. A 300 μL aliquot of splenic homogenate was added to 1 mL Nuclei EZ Lysis Buffer and both the testis and splenic samples were incubated for 5 minutes on ice, with mixing by gentle inversion after 3 minutes. To avoid somatic contaminant, 600 μL of the testis suspension was transferred to a fresh 1.5 mL Lo- Bind microcentrifuge tube. Suspensions were centrifuged at 500 ***×*** g for 5 minutes at 4°C. The supernatant was removed, and the pellet was resuspended with 1 mL of Nuclei EZ Lysis Buffer. After a 5 minute incubation on ice, the samples were again centrifuged at 500 g for 5 minutes at 4°C. Without disrupting the pellet, 1 mL of Nuclei Wash and Resuspension Buffer (NWRB; DPBS with 1% [v/v] BSA) was slowly added and allowed to buffer exchange for 5 minutes on ice. The pallet was resuspended by gentle pipetting up and down 10 times using a 1 mL wide- bore pipette tip. Samples were centrifuged at 500 ***×*** g for 5 minutes at 4°C, and most of the supernatant was removed, leaving behind just enough to cover the pellet. Each sample was resuspended with 300 μL NWRB supplemented with DAPI (10 μg / μL), and then filtered using a 40 μm Flowmi Cell Strainer (Bel-Art) into a 5 mL polypropylene tube. Samples were FACS sorted and haploid nuclei (peaks with DAPI content half of diploid control peaks) were collected.

### Bulk ATAC-sequencing of haploid sperm nuclei

Bulk tagmentation was performed using either commercial pre-assembled Tn5 transposase (Diagenode) or with home-made Tn5 transposases produced and assembled using previously published methods (Picelli et al., 2014) (Supplementary figure 1).

Libraries were prepared using previously published methods (Corces et al., 2017) with minor modifications. Approximately 50,000 haploid sperm nuclei were FACS sorted into a 5 mL polypropylene tube containing 300 uL NWRB with 0.1% (w/v) BSA. Nuclei were centrifuged at 500 ***×*** g for 5 minutes at 4°C and the supernatant was aspirated. Without disrupting the pellet, 1 mL of OMNI-ATAC RSB buffer (10 mM Tris–HCl pH 8.0, 10 mM NaCl, 3 mM MgCl2) was slowly added and the sample was incubated for 5 minutes, then resuspended gently using a wide bore tip. Nuclei were centrifuged at 500 ***×*** g for 5 minutes at 4°C and the supernatant was aspirated. Nuclei were resuspended in 100 μL chilled OMNI-ATAC RSB buffer containing 0.1% (v/v) Tween 20, 0.1% (v/v) IGEPAL, and 0.01% (v/v) Digitonin and incubated on ice for 2 minutes. Nuclei were centrifuged again at 500 ***×*** g for 5 minutes at 4°C and the supernatant was aspirated. Nuclei were resuspended in 50 μL freshly prepared transposition mix (33 mM Tris–HCl pH 8.0, 66 mM Potassium Acetate, 10 mM Magnesium Acetate, 15% (v/v) N,N-dimethylformamide, 0.01% (v/v) Digitonin, 2.5 μL Tn5 assembled enzyme) and mixed by pipetting up and down ten times. Transposition reactions were incubated at 37°C for 45 minutes in a thermocycler with gentle mixing by tapping the tube at 5-minute intervals. Transposition reaction was stopped with 50 μL 2xATAC-STOP buffer (10 mM Tris–HCl pH 8.0, 20 mM EDTA). Afterwards, the samples were purified using a Monarch Nuclei Acid Purification Kit (NEB) as per manufacturer’s instructions and eluted in 20 μL of nuclease free water. All libraries were amplified with barcoded primers listed below, using previously described methods (Buenrostro et al., 2015).

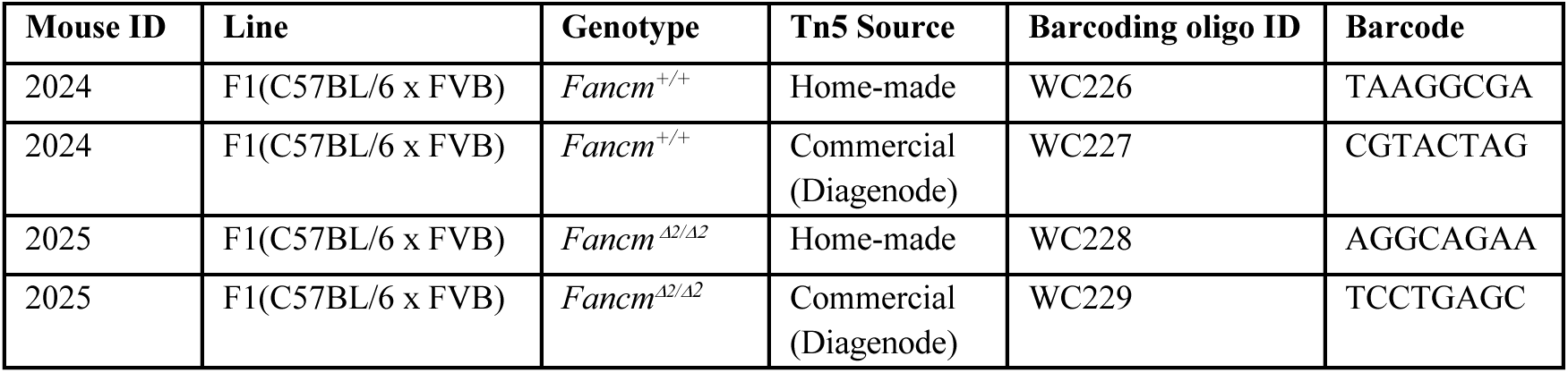

### Single haploid sperm ATAC-seq library preparation and sequencing

Preparation of single-gamete ATAC-sequencing libraries was performed with modifications to our previously published protocol (Novakovic et al., 2022) (ref SSNIP-seq paper DOI: 10.1371/journal.pone.0275168). Briefly, for each single-gamete ATAC-sequencing library, haploid sperm nuclei were FACS sorted into a 5 mL polypropylene tube containing 300 µL NWRB with 0.1% (w/v) BSA. Nuclei were centrifuged at 500 ***×*** g for 5 minutes at 4°C and the supernatant was aspirated. Nuclei were resuspended in NWRB with 0.1% (w/v) BSA supplemented with 0.01% (v/v) Digitonin and incubated on ice for 5 minutes. Nuclei were centrifuged again at 500 ***×*** g for 5 minutes at 4°C and the supernatant was aspirated. 500 μL of Diluted Nuclei Buffer (10x Genomics) was slowly added to the nuclei without disrupting the pellet and allowed to buffer exchange for 5 minutes on ice. The nuclei were resuspended in 10 μL of Diluted Nuclei Buffer (10x Genomics) and quantified using a hemocytometer. Nuclei concentration was adjusted to attain a capture number of around 1,000 nuclei, accounting for loss during subsequent steps. Prepared nuclei were added to the Transposition Mix (10x Genomics) and transposition, library preparation and clean-up for single-cell ATAC-sequencing was performed according to the manufacturers protocol using the Chromium Single-Cell ATAC Kit v1.1 (10x Genomics). Each library was sequenced on the NovaSeq 6000 (Illumina) using a 50/8/16/50 bp read configuration.

### Data processing

Bulk ATAC-sequencing fastq files were trimmed using *cutadapt* (v3.4) (Martin, 2011) and mapped to the mm39 reference genome using *bwa-mem* with default settings (H. Li, 2013). Duplicate reads were removed using MarkDuplicates (*picard tools*).

For snATAC-sequencing, fastq files were processed using Cell Ranger ATAC (v2.1.0; 10x Genomics) to align reads to the mm39 reference genome. Outputs from Cell Ranger were used in downstream analysis.

### Sequencing coverage analysis

Coverage files for bulk and snATAC experiments were generated from BAM files using

*samtools depth* (H. Li et al., 2009). Data were summarised in 10 kb bins based on chromosome

lengths from mm39. Genomic tracks were visualised with *Gviz*, with gene annotations added from UCSC genome browser (Hahne & Ivanek, 2016).

### Single nuclei ATAC-sequencing analysis with *Signac* and *Seurat*

The outputs from Cell Ranger containing peak and fragment information were used to generate coverage estimates of the snATAC-seq data by identifying a set of common peaks across all samples, indicative of open chromatin regions. Barcodes with fewer than 500 fragments per cell were filtered out. Fragment counts were then normalised using term-frequency inverse-document-frequency (TF-IDF) to result in the logarithmic ratio of the total number of reads per cell, relative to total cells.

The normalised data was used to assess coverage in specified genetic regions, as well as to assess various QC metrics, including: TSS enrichment scores, fragment counts (nCounts), nucleosome signal, and percentage of reads in peaks. Dimensionality reduction was performed using Singular Value Decomposition (SVD), and Uniform Manifold Approximation and Projection (UMAP) plots were generated to visualise clusters, with Latent Semantic Indexing (LSI) applied for data reduction. Spare clusters were filtered, and UMAP plots were separated into sequencing batches. Gene activities were annotated using the Ensembl database and the mm39 reference genome. Cluster-specific markers with positive fold changes between clusters were identified, and *Enrichr* was used to identify putative cell types based on the top differentially accessible genes.

### Single nuclei ATAC-sequencing crossover calling with sgcocaller and comapr

Crossover calling was conducted on the Cell Ranger BAM output file using *sgcocaller* (Lyu et al., 2022), with the following parameters: --cmPmb 0.1 --maxDP 20 --maxTotalDP 450 --

minTotalDP 0 --minDP 0 --thetaREF 0.2 --thetaALT 0.8. FVB/N-specific variants differing from the mm10 reference genome were obtained from the Mouse Genome Project (FVB_NJ.mgp.v5.snps.dbSNP142.vcf) and lifted over for compatibility with the mm39 reference genome.

The outputs from Cell Ranger were filtered to generate a list of barcodes from intact cells with greater than 10,000 reads. To refine crossover calls, the filtered barcode file and output of *sgcocaller* were processed utilising *comapr* (Lyu et al., 2022), generating genetic distance maps with the Kosambi mapping function, and identifying inter-crossover distances. The following thresholds were set in *comapr*: minSNP = 10, minCellSNP = 100, maxRawCO = 5, minLogllRatio = 30, bpDist = 1e5. Artefacts with double crossovers occurring within 3 bp were removed from the analysis.

### Marker segregation for single nuclei ATAC-sequencing

Segregation states of SNPs were analysed in 10 Mb bins, based on chromosome lengths from the mm39 reference genome, using output files generated by *sgcocaller*. SNP positions and their underlying segregation states were used to calculate the ratio of BL6 and FVB/N genetics, where 1 indicated FVB/N and 0 indicated BL6. Haplotype ratios were computed, and binomial distribution tests were performed to assessed bias in marker segregation. Telomeric regions were removed due to low coverage, which resulted in inaccurate state imputation.

### Permutation tests and null distributions generation

Permutation tests using label swapping and null distribution generation were conducted following previously described methods (Tsui et al., 2023).

### ChIP-seq compared to CO analysis

C57BL/6 PRDM9 and DMC1 ChIP-seq data (Biot et al., 2024) were downloaded from NCBI’s Sequence Read Archive (SRA), using the SRA toolkit. ChIP-seq reads were aligned to the mm39 reference genome using *bowtie2* (Langmead & Salzberg, 2012) and filtered to remove low-quality reads and duplicates using *samtools* (fixmate, sort, and markdup; (H. Li et al., 2009)). PRDM9 and DMC1 knockout samples were excluded from wild-type analysis, and ChIP-seq peaks were called using MACS2 (Zhang et al., 2008).

CO positions were derived from the snATAC-seq data, processed using *sgcocaller* and *comapr*. Overlapping and ChIP-seq and CO positions were identified using the *findOverlaps* function from *GenomicRanges* (Lawrence et al., 2013). Statistical comparisons were conducted using one-sided Wilcoxon signed-rank tests, and as permutation tests, comparing observed ChIP-seq signals to the null distribution.

## Oligonucleotide sequences used in this study

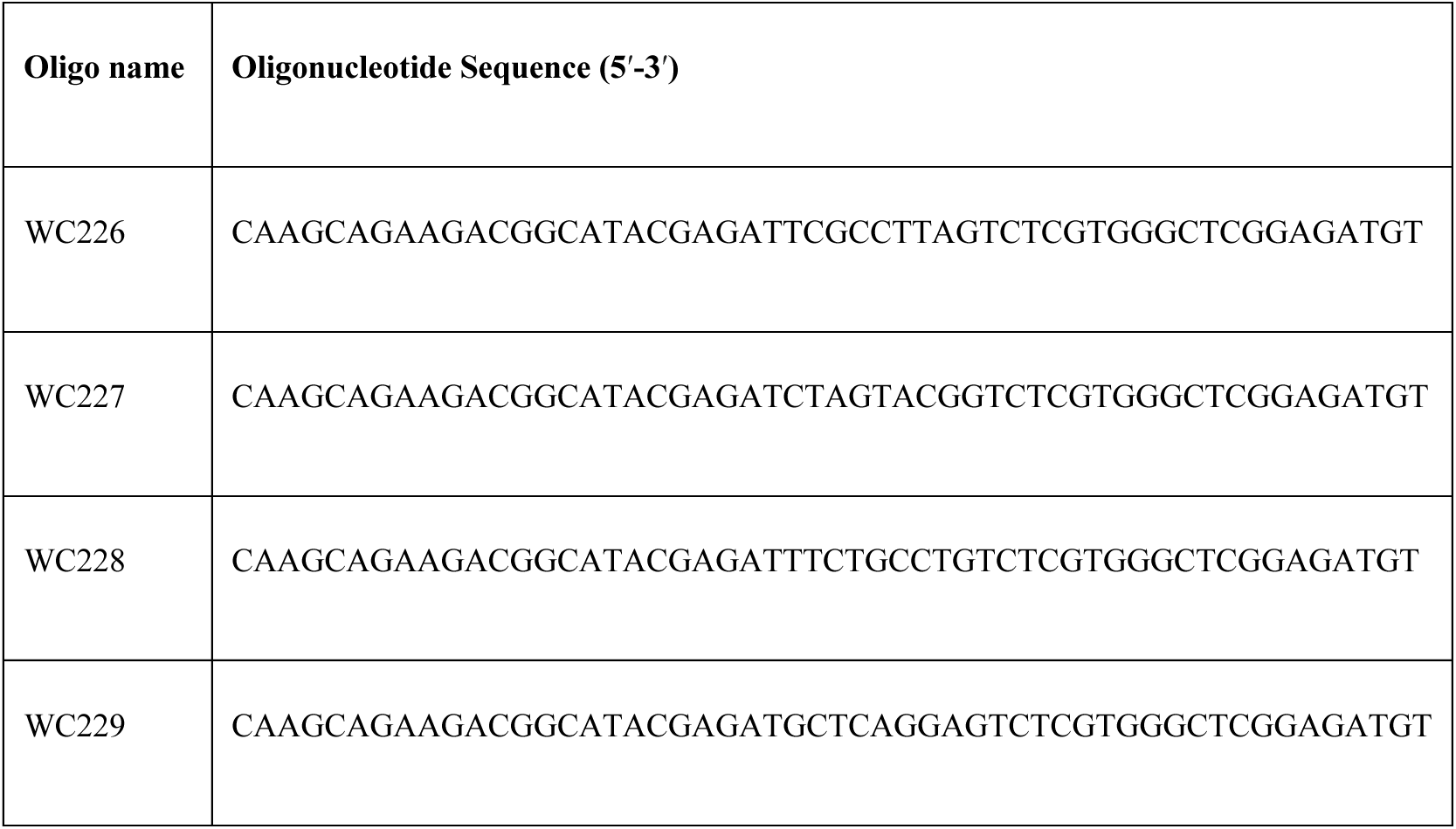

---

**Supplementary Figure 1.**
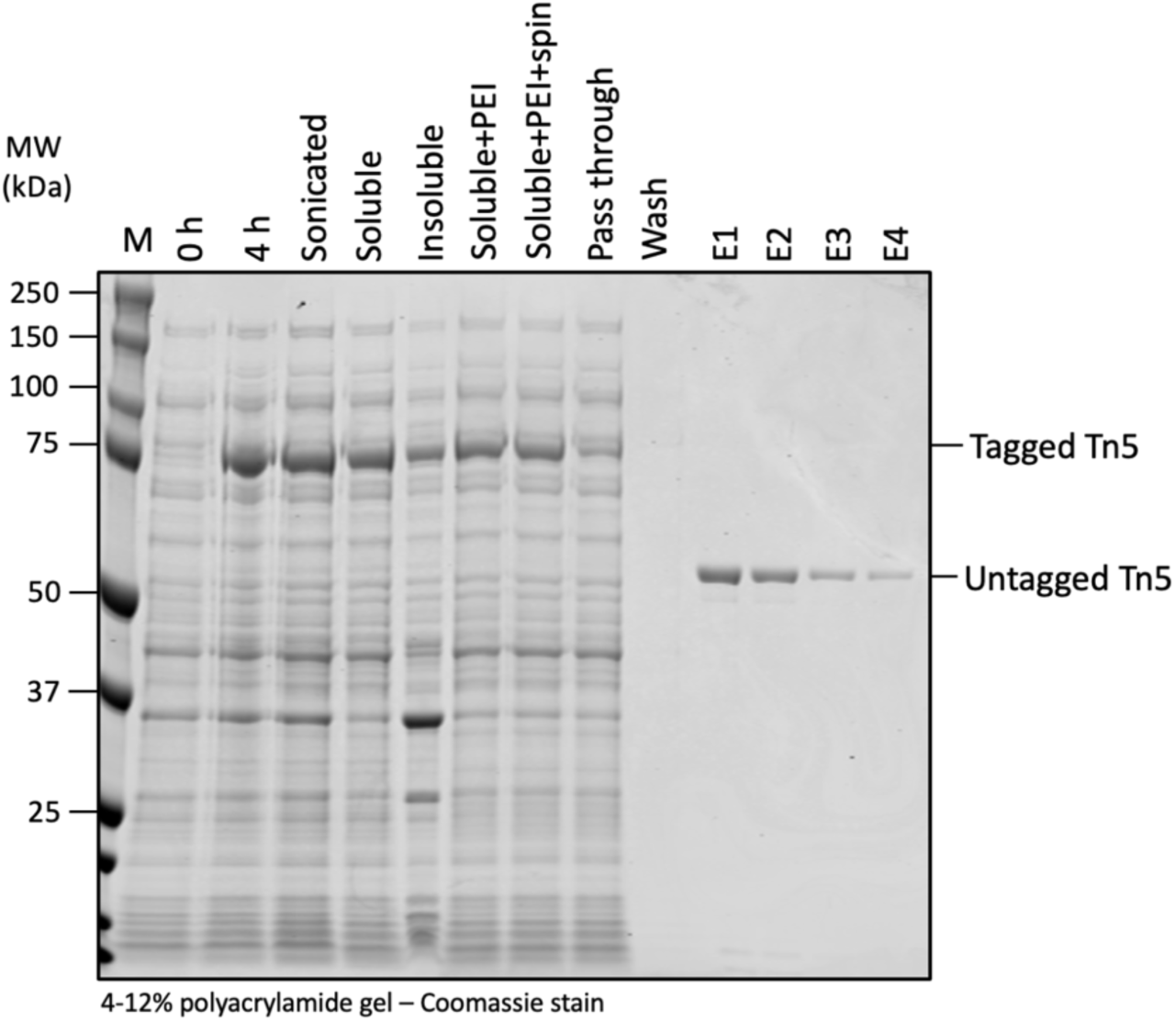
SDS-PAGE analysis of Tn5-intein construct expression and purification. Samples were collected at various stages of expression at 23°C and purification process and assessed by SDS-PAGE to evaluate the presence and purity of the Tn5-intein protein. Analysed samples include: molecular weight marker; crude lysate at 0 hours (0 h) and 4 hours (4 h) post-induction with IPTG; cell pellet post-sonication (Sonicated); the soluble (Soluble) and insoluble (Insoluble) fraction post-sonication; the soluble fraction treated with polyethyleneimine (soluble+PEI); the soluble fraction treated with PEI and then centrifuged (Soluble+PEI+spin); the Flow-through after loading lysate onto a chitin column (pass through); the wash fraction from chitin column (Wash); the eluted fractions (E1 to E4) from the chitin column. Molecular weight (MW) of the marker in kilodaltons (kDa) shown on the left. The expected size of tagged and cleaved (untagged) Tn5 is indicated on the right.

**Supplementary Figure 2.**
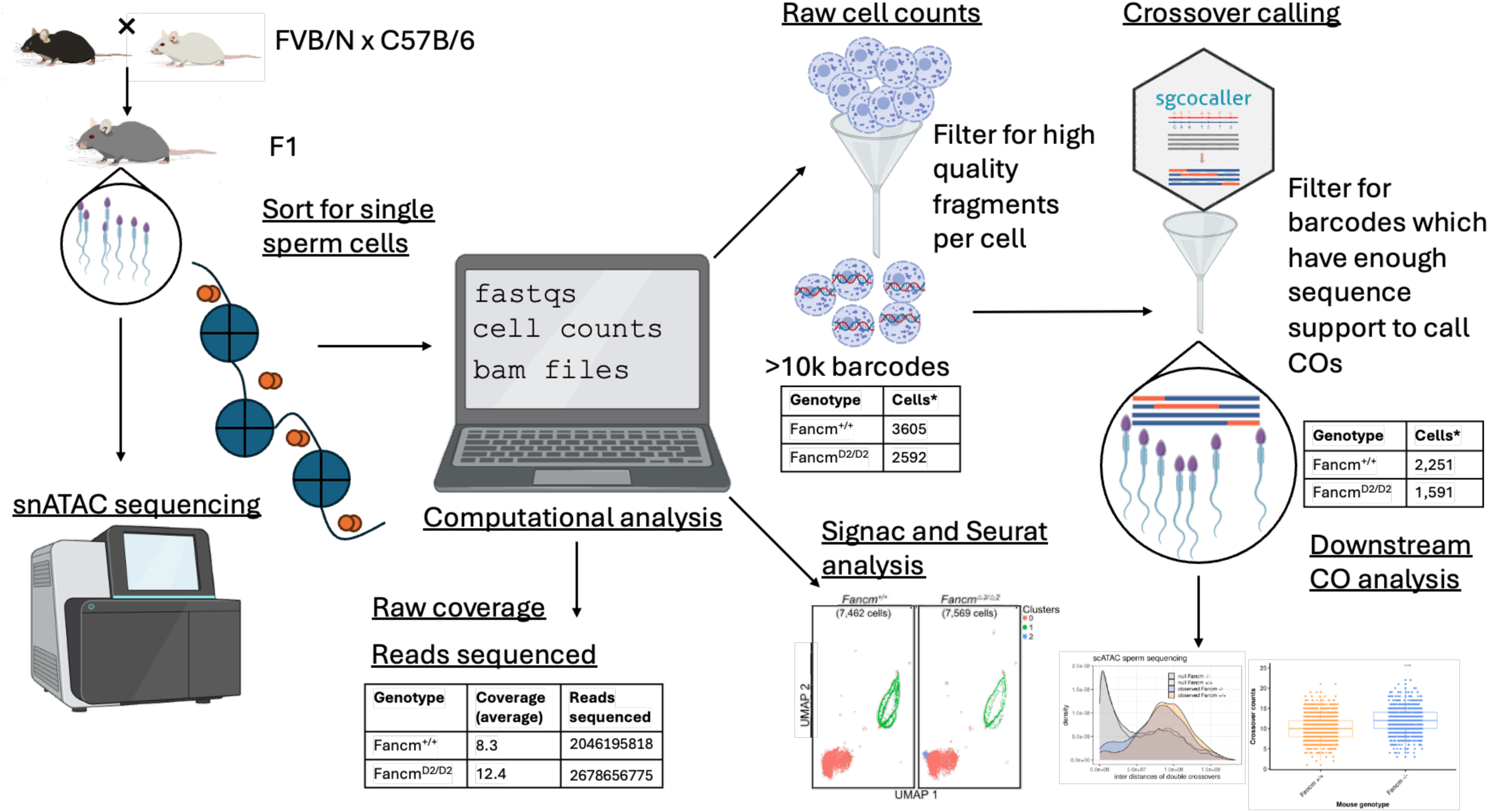
Workflow of snATAC computational analysis. Nuclei were extracted and sequenced from F1.*Fancm^+/+^* and F1.*Fancm^Δ2/Δ2^*mice, generated by crossing FVB/N and C57BL/6 inbred strains. Fastq files from ATAC sequencing were processed using Cell Ranger ATAC (10x Genomics) to generate read counts, cell counts, and coverage metric. Peak files were utilised in Seurat for downstream analysis, including filtering and processing to create UMAP plots. Crossover analysis was performed using BAM files output by Cell Ranger ATAC, which were filtered to retain cells over 10,000 barcodes. Crossover calling was conducted on the filtered samples using *sgcocaller*, followed by additional filtering with *comapr* to ensure sufficient support for crossover identification. These filtered datasets were then used for downstream analysis.

**Supplementary table 1:** Summary of sequencing quality and crossovers detected in snATAC sequencing.

| Sample ID | Genotype | Unfiltered cells | High-quality fragments, >10k per barcode | High-quality fragments per cell, >10k per barcode | Total number of sequenced read pairs | Average coverage per BAM file | Barcodes passed filtering for COs - number of sperm | Mean COs per sample | Single COs | Double COs | Triple COs | Quadruple COs |
| --- | --- | --- | --- | --- | --- | --- | --- | --- | --- | --- | --- | --- |
| 1826 | *Fancm*<sup>+/+</sup> | 1,007 | 687 | 579 | 881,941,291 | 10.3488 | 173 | 10.7 | 1365 | 133 | 13 | 0 |
| 1919 | *Fancm*<sup>+/+</sup> | 1,674 | 1794 | 1664 | 555,467,003 | 9.42015 | 1200 | 11.1 | 7796 | 875 | 31 | 4 |
| 1929 | *Fancm*<sup>+/+</sup> | 889 | 1417 | 1362 | 608,787,524 | 5.05501 | 878 | 9.7 | 5617 | 394 | 14 | 1 |
| 1825 | *Fancm*<sup>Δ2/Δ2</sup> | 1,452 | 1083 | 923 | 1,015,085,960 | 18.5083 | 412 | 11.6 | 2678 | 366 | 26 | 7 |
| 1921 | *Fancm*<sup>Δ2/Δ2</sup> | 785 | 833 | 783 | 585,535,532 | 8.6143 | 539 | 12.2 | 3683 | 563 | 31 | 2 |
| 1928 | *Fancm*<sup>Δ2/Δ2</sup> | 889 | 940 | 886 | 1,078,035,283 | 9.96556 | 640 | 12.1 | 4135 | 688 | 36 | 2 |

**Supplementary table 2:**
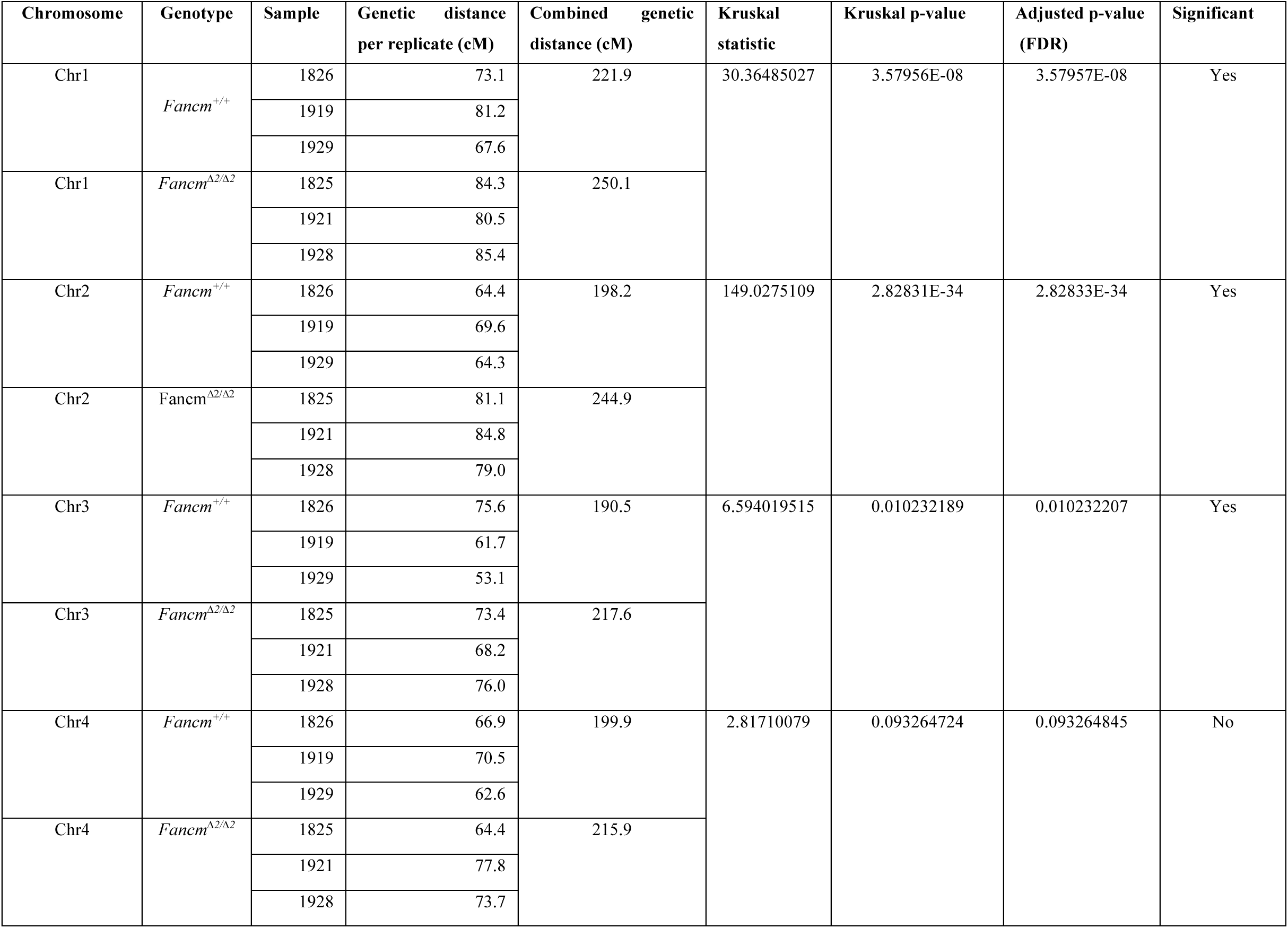
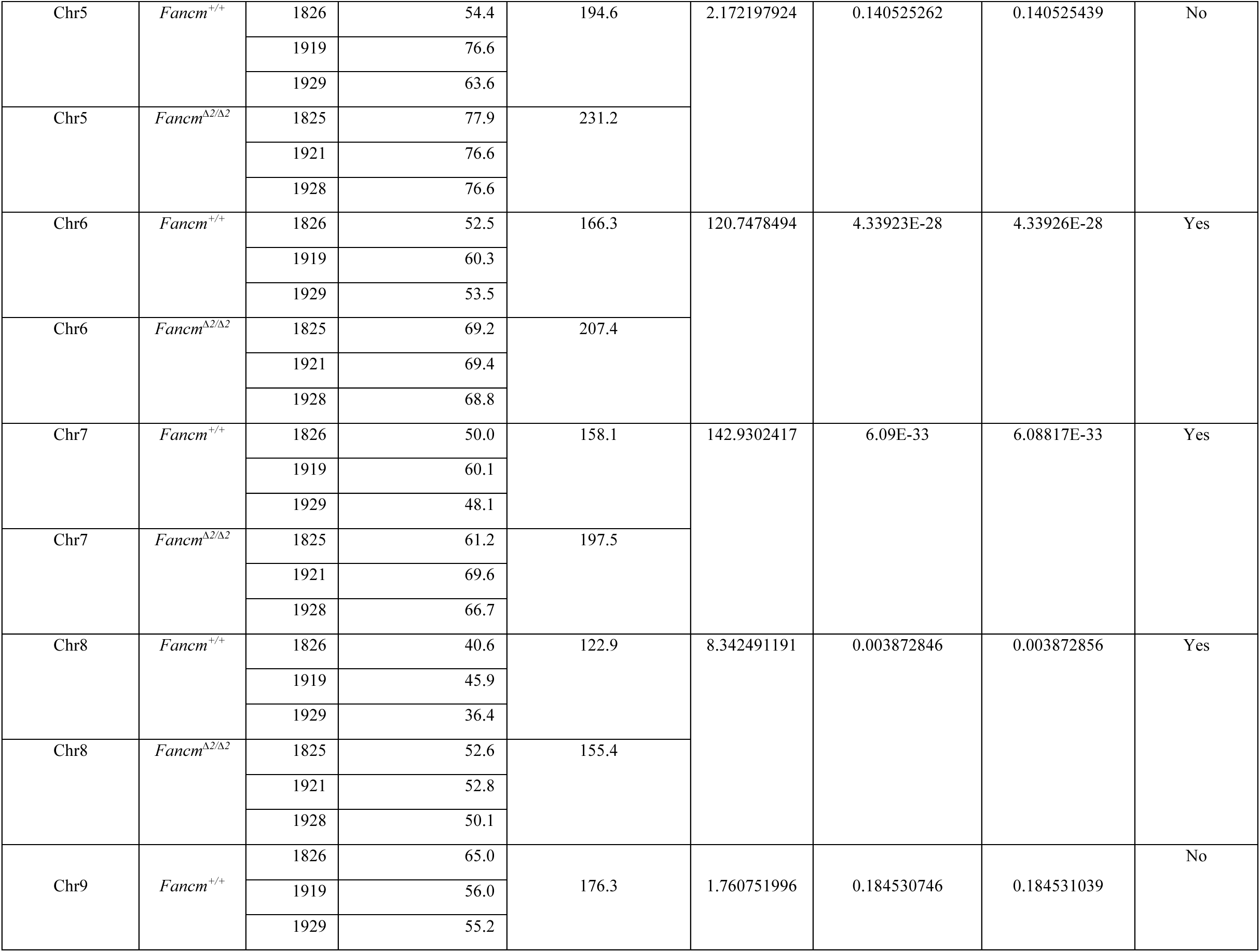
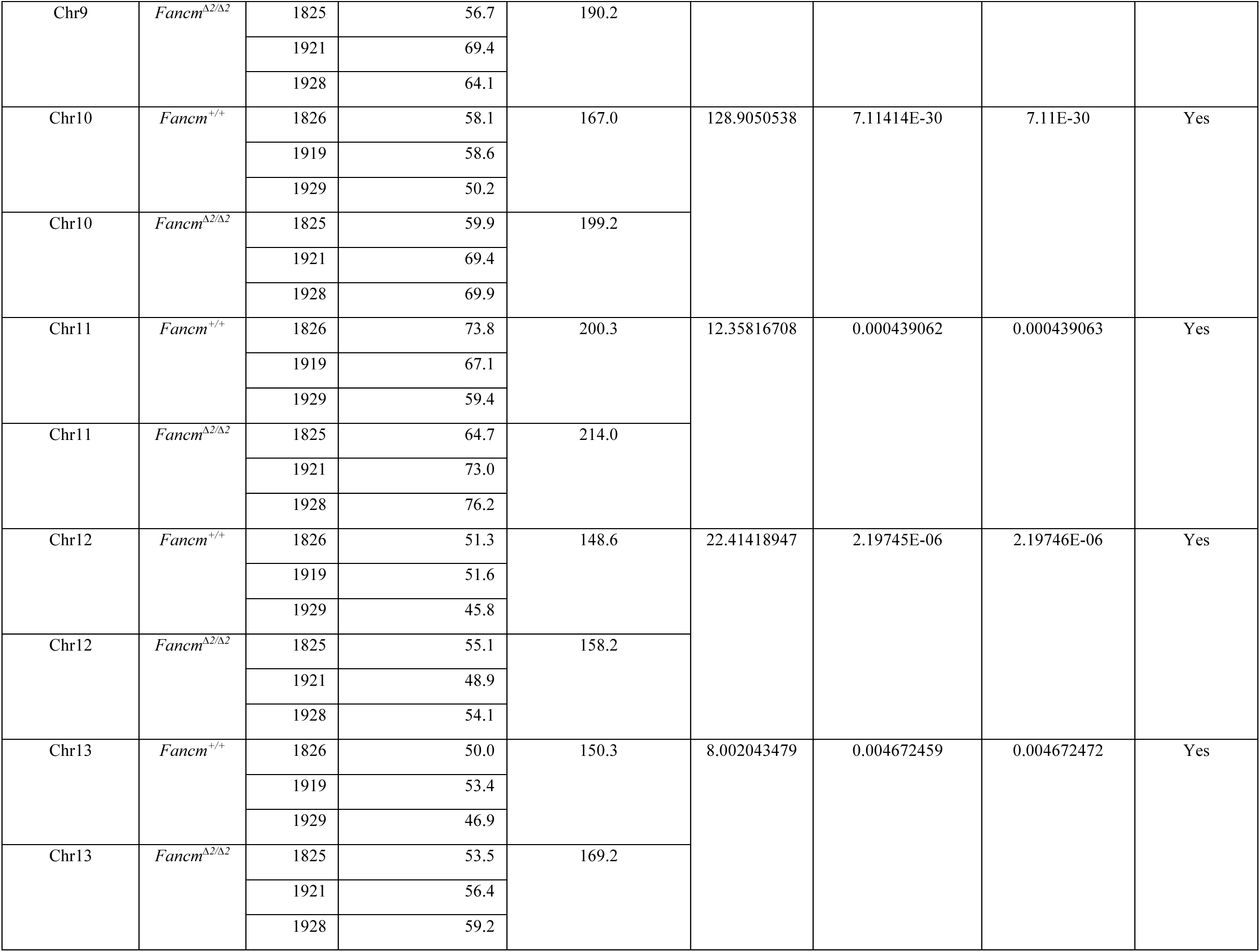
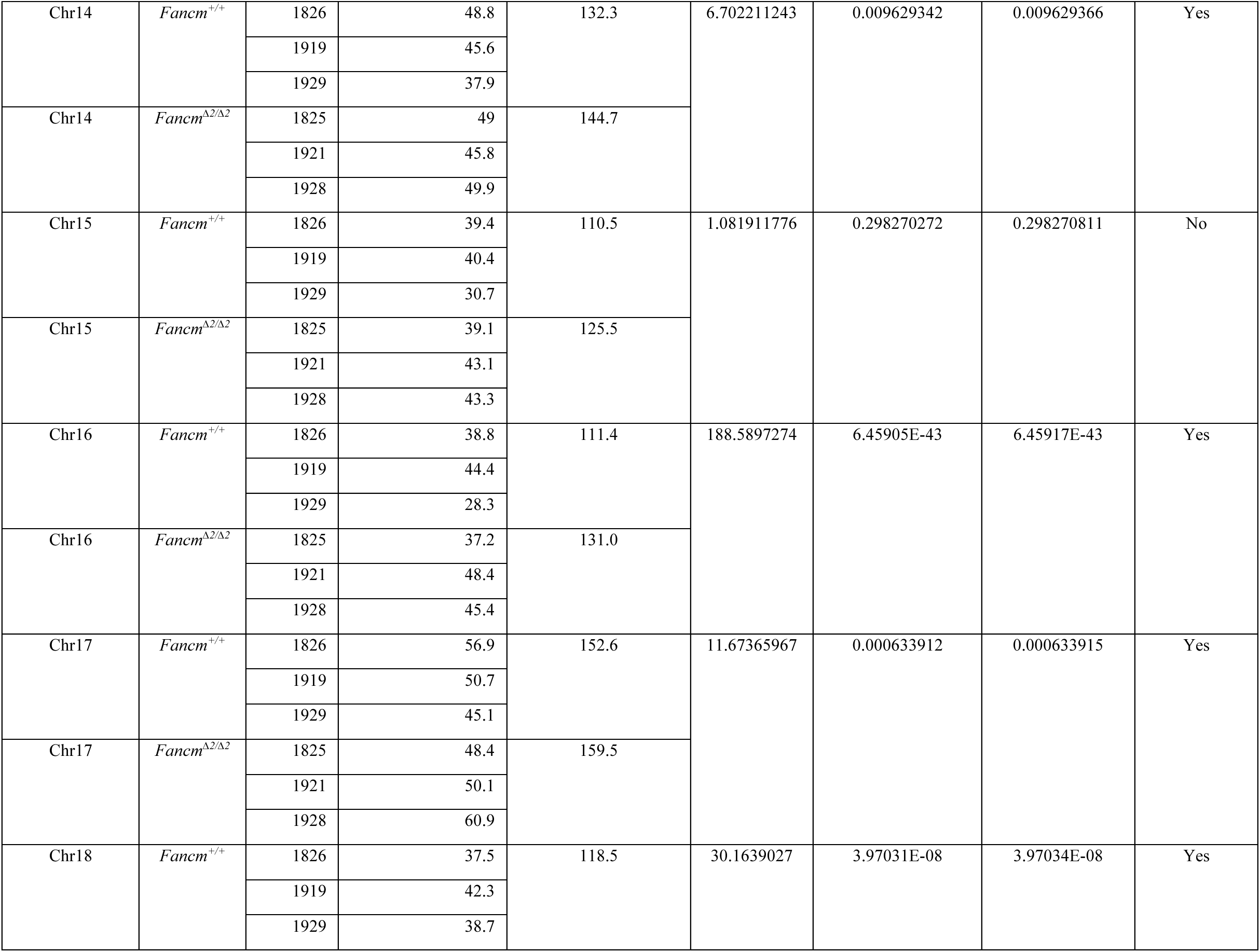
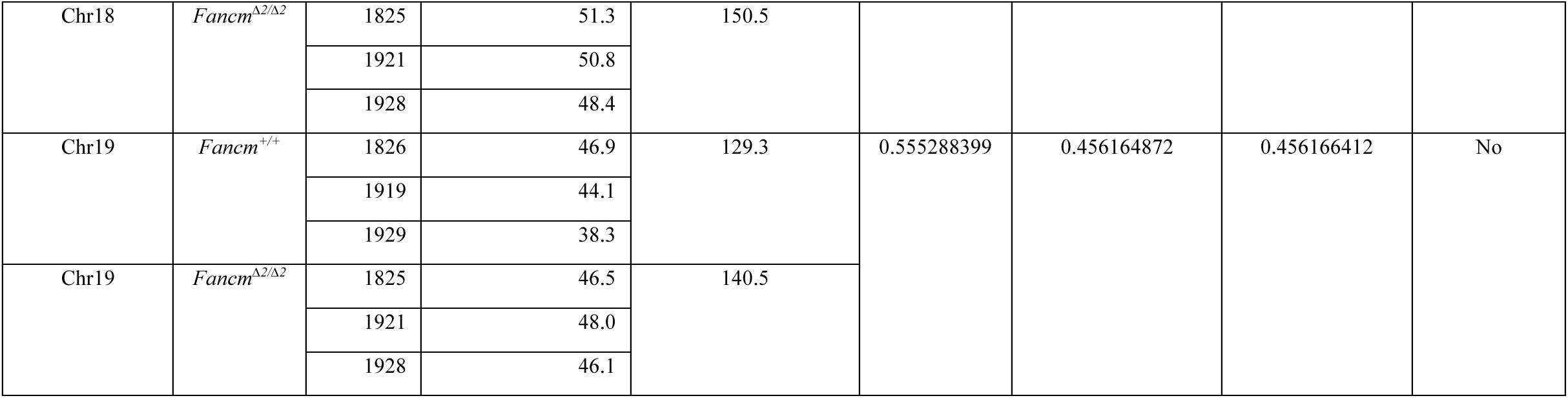
Genetic distances of each chromosome with a comparison between genotypes, with p corrected p-values.

